# RNA LEVER Mediates Long-Range Regulation of ε-globin by Keeping PRC2 in Check

**DOI:** 10.1101/2020.09.05.282624

**Authors:** Wei Wen Teo, Xinang Cao, Chan-Shuo Wu, Hong Kee Tan, Qiling Zhou, Henry Yang, Li Chai, Daniel G. Tenen

## Abstract

Polycomb Repressive Complex 2 (PRC2) is an epigenetic regulator required for gene silencing during embryonic development. Previous studies have reported that PRC2 interacts with RNA in a promiscuous manner, but the biological functions of such interaction are unknown. Here we present a seesaw mechanism for the regu<u>**l**</u>ation of ε-globin through inacti<u>**v**</u>ating <u>**E**</u>ZH2 by an upstream non-coding <u>**R**</u>NA (LEVER). We show that LEVER, a non-coding RNA identified by RNA immunoprecipitation sequencing (RIP-seq) of the PRC2 core subunit EZH2 and Nanopore sequencing, binds PRC2 and thereby prevents the accumulation of H3K27 methylation along the genomic region where LEVER RNA is transcribed. The open chromatin within the LEVER locus in turn competes for the chromatin interaction between the ε-globin promoter and the Locus Control Region (LCR), working as a negative regulatory element of ε-globin expression. Hence, LEVER RNA negatively regulates ε-globin by sequestering PRC2 from repressing the LEVER locus, which is a competitor of the ε-globin-LCR interaction.

## Introduction

Polycomb Repressive Complex 2 (PRC2) is an epigenetic regulator that plays essential roles in cellular differentiation and early embryonic development. These biological functions are achieved mainly through establishing and maintaining the global methylation landscape of lysine 27 on histone H3. In particular, the final H3K27 tri-methylated product (H3K27me3) generally marks transcriptionally inactive regions and is considered as the major epigenetic output through which PRC2 supervises the repressive genome ^1^.

However, in mammals, how PRC2 selectively represses targets globally remains elusive. Many long non-coding RNAs (lncRNAs), such as XIST ^2^ and HOTAIR ^3, 4^, have been proposed to serve as co-factors to direct PRC2 to its specific targets in the mammalian genome. Importantly, this model highlights a specific lncRNA-PRC2-genome loci axis.

This lncRNA-directed PRC2-mediated gene repression model has been questioned by several follow-up studies on PRC2-RNA interaction. First, the promiscuous RNA binding pattern of PRC2 *in vitro* and in cells challenges the selectivity of PRC2 for its specific lncRNA partners ^5^. Second, RNA can inhibit PRC2 enzymatic activity ^6^ through competing with histone for PRC2 binding *in vitro*^7^. Furthermore, extensive interaction between PRC2 and nascent RNAs was observed in mouse embryonic stem cells (ESCs) at transcriptionally active gene loci ^8, 9^, which brings into question the transcription repressive effect by local PRC2-RNA interaction at these loci. In line with these findings, blocking RNA transcription in mouse ESCs increases global PRC2 occupancy and H3K27me3 levels at previously active gene loci ^10^.

To investigate the biological functions of PRC2-RNA interactions, we identified robust PRC2-bound RNAs through RIP-seq in K562 cells. This led to the identification of a novel PRC2-interacting non-coding RNA (termed LEVER, short for the regu<u>**l**</u>ation of ε-globin through inacti<u>**v**</u>ating <u>**E**</u>ZH2 by an upstream non-coding **R**NA), and we further characterized LEVER by direct-RNA Nanopore sequencing. Depletion of LEVER RNA increased PRC2 occupancy and H3K27me2/me3 levels at the LEVER locus. Importantly, down-regulation of LEVER enhanced the expression of the downstream neighboring ε-globin gene. Mechanistically, the epigenetic change at the LEVER locus caused by increased PRC2 binding upon LEVER RNA depletion remodeled the local chromatin interaction involving the β-globin Locus Control Region (LCR) and thereby activated ε-globin transcription. In summary, our study reveals a model that LEVER RNA sequesters PRC2 from its genome locus and blocks PRC2 activity *in cis*, which remodels local chromatin interaction and regulates neighboring gene expression.

## Results

### EZH2 binds G/A-rich nascent transcripts

To identify PRC2 associated RNAs, we performed RIP-seq in K562 cells for endogenous EZH2, which is the enzymatic subunit and the subunit with the highest RNA-binding affinity in PRC2 ^6^. The two well-established PRC2-interacting lncRNAs, XIST ^2^ and KCNQ1OT1^*11*^, were both consistently enriched in our biological replicates (Supplementary Figure 1A and 1B), validating our EZH2 RIP-seq. 1528 high-confidence EZH2-interacting RNA fragments were reproducibly identified from both replicates using a stringent peak-calling algorithm. Annotation of these fragments indicates no preference for transcripts generated from coding genes, non-coding genes, or intergenic regions (Figure 1A). Detailed feature annotation of the fragments from the annotated gene loci further reveals that EZH2 preferentially binds intronic rather than exonic RNAs, disregarding the coding potential of the corresponding gene (Figure 1A). To further study the identity of these EZH2-interacting RNAs, we examined their cellular localization by quantifying their reads-per-million-reads (RPM) values in RNA-seq of nuclear or cytoplasmic fractions of K562 cells. 89% of these EZH2-bound RNA fragments have higher read counts in nuclear than in the cytoplasmic fraction, suggesting their nuclear localization (Figure 1B). Further comparison of the RNA-seq read counts between nuclear polyA+ and nuclear polyA-fractions suggests more than 75% of these EZH2-interacting fragments are non-poly-adenylated (Figure 1B). Collectively, these results support previous findings ^9^ that EZH2 predominantly binds nascent transcripts in the nuclei.

**Figure 1.**
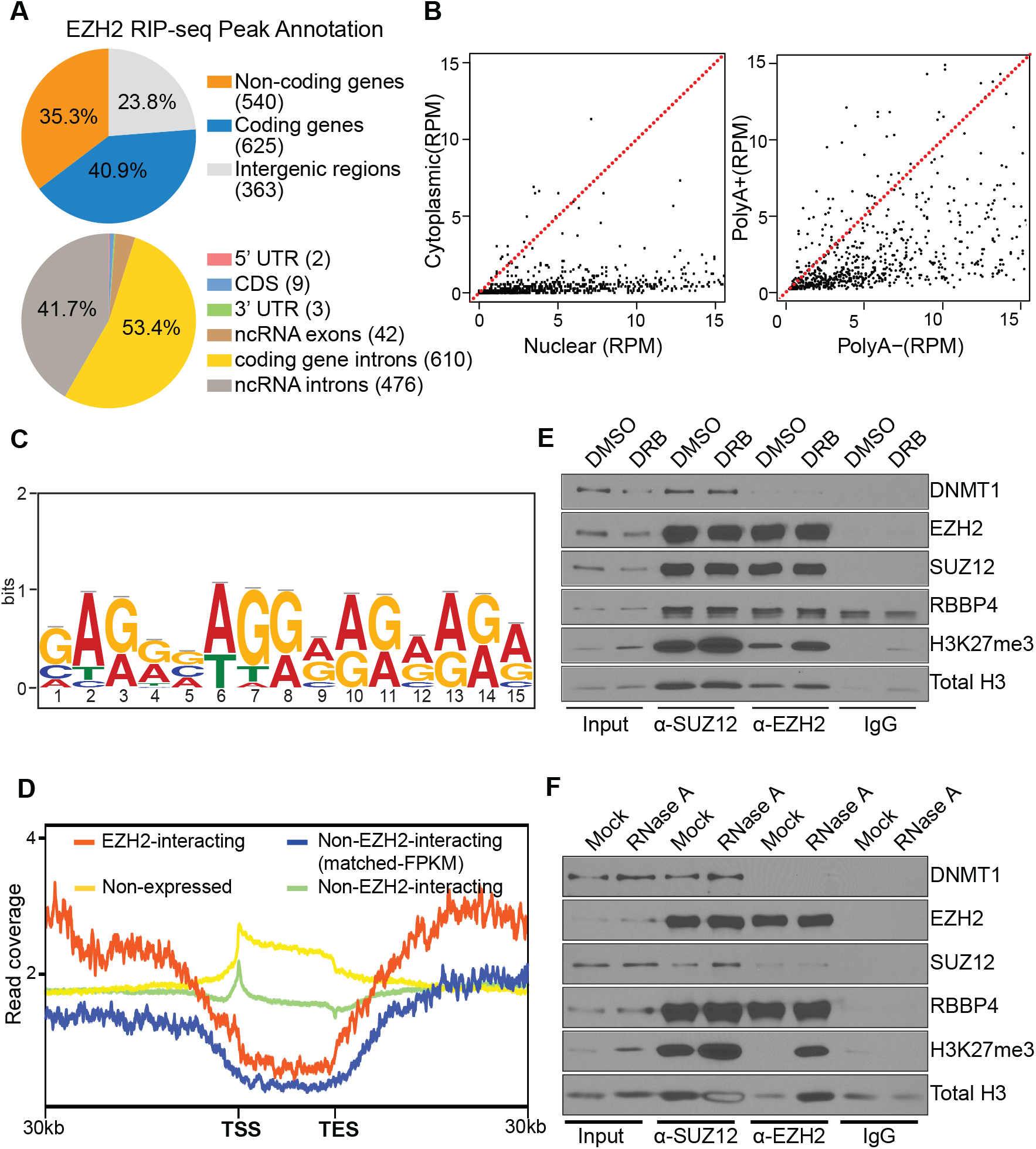
Nascent RNA binds and sequesters EZH2 from histone H3 in K562 cells. (A) EZH2 preferentially binds intronic rather than exonic RNAs. **Upper panel**: Percentage of EZH2 RIP-seq enriched RNA fragments annotated to coding genes, non-coding genes, or intergenic regions. The number of identified peaks annotated to each category are labelled in the brackets in the pie-chart legends. **Lower panel**: Percentage of EZH2 RIP-seq enriched RNA fragments originated from coding or non-coding genes annotated to different gene features. UTR, untranslated regions; CDS, coding sequences; ncRNA, non-coding RNA. (B) EZH2 predominantly binds nuclear polyA-nascent transcripts. **Left panel**: Scatter plot showing normalized read counts in nuclear (x-axis) and cytoplasmic (y-axis) fractioned RNA-seq for each EZH2 RIP-seq enriched RNA fragment. The read counts are normalized to total mapped reads as RPM (reads per million mapped reads). Enrichment of dots below the labeled diagonal reveals higher RNA-seq reads in the nuclear than in the cytoplasmic RNA fraction for most of the EZH2 RIP-seq fragments. **Right panel**: similar as in left, except that the read counts of the identified RNA fragments are quantified in the nuclear polyA-(x-axis) and nuclear polyA+ (y-axis) fractions. (C) RNA motif identified from a set of 900 EZH2-bound RNA fragments by MeMe motif discovery analysis. (D) Read coverage profiles for H3K27me3 ChIP-seq spanning the upstream, gene body, and downstream regions of the following gene sets in K562 cells: Red line: EZH2-interacting genes; blue line: non-EZH2-interacting genes with matched expression level as EZH2-interacting genes; green line: total non-EZH2 interacting expressed genes; and yellow line: non-expressed genes. Notably, the “EZH2-interacting” stated here is based on the RNA-binding as identified by EZH2 RIP-seq rather than ChIP-seq (chromatin binding) of EZH2 at these gene loci. Shown in the X-axis is distance from the transcription start site (TSS) to the transcription end site (TES). (E) (F) EZH2 or SUZ12 co-immunoprecipitation (co-IP) in K562 cells treated with (E) 5,6-dichloro-1-β-D-ribofuranosylbenzimidazole (DRB) or (F) RNase A. Co-IP of PRC2 subunits (EZH2, SUZ12, RBBP4), PRC2-interacting protein (DNMT1), and histones (total histone H3 and histone H3 with H3K27me3 methylation) were measured by western blot. IgG isotype control antibody was used as co-IP negative control.

To further investigate whether EZH2 possesses any binding preference for specific RNA sequences, we performed motif discovery analysis for EZH2-enriched RNA fragments with MeMe Suite ^12^. Two sets of flanking sequences were included in the analysis to eliminate background noise. One motif rich in guanidine (G) and adenosine (A) was significantly enriched from our motif analysis (Figure 1C). This G/A-rich motif revealed by RIP-seq in cells highly agrees with previous results from *in vitro* RNA binding experiments ^13^. Collectively, our EZH2 RIP-seq in K562 cells reveals that EZH2 prefers to bind G/A-rich nascent transcripts in cells.

### EZH2-bound nascent transcripts antagonize PRC2 activity

As nascent transcripts are geometrically close to their corresponding gene loci, nascent transcript-EZH2 interaction may locally influence the histone methylation function of PRC2. To investigate this, we profiled genome-wide H3K27 methylation status as the readout of PRC2 activity at the gene loci where EZH2-interacting transcripts are generated (Figure 1D). We refer to these gene loci as “EZH2-interacting gene loci”, while “non-EZH2-interacting gene loci” refer to those in which no EZH2-bound transcript was identified by our RIP-seq. This definition of ‘interacting’ is based on RNA-binding (RIP-seq) instead of the chromatin binding (ChIP-seq) of EZH2. Consistent with the nature of a repressive histone mark, the H3K27me3 level is remarkably higher in the gene body region (from the transcription start site (TSS) to the transcription end site (TES)) of non-expressed genes compared with their flanking regions. In contrast, the H3K27me3 of the EZH2-interacting gene loci showed the opposite pattern, reflecting active transcription in these loci. We further compared the H3K27me3 level of EZH2-interacting gene loci with non-EZH2-interacting gene loci. To minimize the epigenetic bias caused by different gene expression levels, a set of non-EZH2-interacting genes with gene-by-gene matched expression levels as those EHZ2-interacting genes were selected for comparison (this set of genes is termed non-EZH2-interacting matched-FPKM, Supplementary Figure 1C). Interestingly, we noticed the EZH2-interacting genes have higher H3K27me3 basal levels in the flanking regions than their FPKM matched non-EZH2-interacting genes and the genomic background (flanking regions of non-EZH2-interacting and non-expressed genes), suggesting their residence in genomic regions that are generally more repressed and with more stable contact with PRC2. Moreover, the decrease of H3K27me3 levels from the flanking regions to the gene body is steeper in the EZH2-interacting gene loci than in the non-EZH2-interacting gene loci, suggesting that RNA-binding may have played a PRC2-inhibitory function. In sum, these observations suggest that EZH2 interacts with RNA within densely H3K27 methylated genomic regions in which the emerging RNA can be crucial in curtailing the repressive activity of PRC2 at the corresponding gene locus.

To further study how RNA affects PRC2 chromatin occupancy biochemically, we inhibited global transcription by treating K562 cells with the RNA Polymerase II (Pol II) inhibitor 5, 6-dichloro-1-β-D-ribofuranosylbenzimidazole (DRB) in culture and performed co-immunoprecipitation against PRC2 subunits EZH2 or SUZ12 (Supplementary Figure 1D). While lack of RNA did not affect the binding of EZH2 or SUZ12 to other PRC2 subunits (SUZ12/EZH2 and RBBP4), both EZH2 and SUZ12 showed increased interaction with lysine 27 tri-methylated histone H3 (H3K27me3) and, to a lesser extent, with total H3 upon transcription blockage (Figure 1E). In line with this observation, the depletion of RNA by RNase A treatment also markedly enhanced PRC2 interaction with H3 and H3K27me3 (Figure 1F). Of note, both DRB and RNase A treated K562 cells showed a global increase in H3K27me3 but not total H3 in the input samples, revealing that RNA depletion enhanced PRC2 binding to the chromatin and led to increased histone methylation levels, supporting previous reports in mESC ^14^. Collectively, these results strongly support the antagonistic effect of RNA on PRC2 chromatin binding and its methyltransferase activity.

### PRC2 binds a non-coding RNA upstream of β-globin cluster

Our EZH2 RIP-seq enriched many uncharacterized RNAs transcribed from unannotated intergenic regions (Figure 1A). Among these uncharacterized RNAs, we identified one long transcript (which we termed LEVER) whose transcription start site locates 235kb upstream of the β-globin gene cluster on chromosome 11 (Figure 2A). Transcripts from this locus were highly enriched by RIP-seq with multiple significant peaks (labeled as LVR1-4), and this was further validated by RIP-qPCR of three independent biological replicates (Figure 2B). Notably, LEVER RNA is enriched by EZH2 at a level comparable to that of XIST, which strongly suggests the validity of LEVER as an EZH2-interacting RNA.

**Figure 2.**
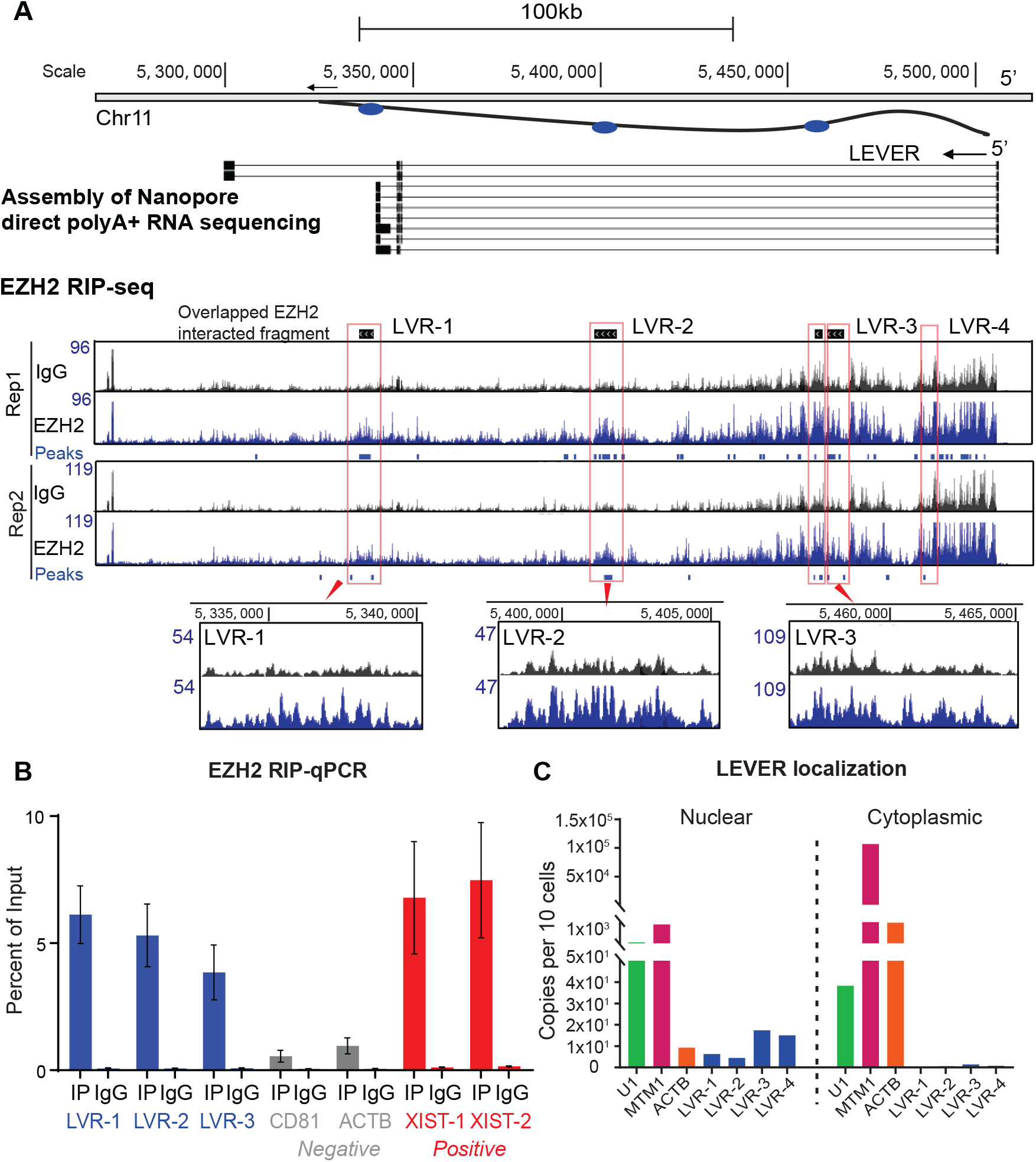
EZH2 binds a nascent transcript at LEVER locus. (A) EZH2 RIP-seq and Nanopore direct sequencing of total polyA+ RNA in K562 cells demonstrating EZH2 binding intronic RNAs from the LEVER locus (pictured in supplementary Figure 2). In the Nanopore track, the bars represent genomic regions covered by actual Nanopore reads, and they are linked by lines to represent splice junctions. In the EZH2 RIP-seq tracks, the EZH2 enriched RNA peaks (blue bars below signal tracks of each replicate) are called by ≥ 2-fold read coverage of EZH2 over IgG with Poisson distribution *p*-value ≤ 10^−5^. Peaks generated from two biological replicates are further overlapped and merged as “Overlapped EZH2 interacted fragments” (black bars above the RIP-seq tracks). Selected overlapped fragments are highlighted in red boxes with their readcoverage tracks shown in a zoomed-in view. The blue ovals in the top diagram represent the relative positions of LVR-1, LVR-2, and LVR-3. (B) EZH2 RIP-qPCR on three LEVER fragments (LVR-1, LVR-2, and LVR-3, highlighted in Figure 2A), CD81 and ACTB (RIP-negative controls) and two XIST fragments (RIP-positive controls), quantified as percent of total input. Data from three biological replicates are shown. (C) Digital droplet RT-PCR of LEVER fragments (LVR-1 through LVR-4), U1 and MTM1 (nuclear enriched), and ACTB (cytoplasmic enriched) RNAs in K562 nuclear or cytoplasmic RNA fractions.

To characterize the nascent and spliced transcripts from the LEVER locus, we assembled the RNA sequences produced by long-read Nanopore cDNA sequencing of K562 nuclear RNA, as well as Nanopore direct sequencing of K562 total polyA+ RNA, which revealed spliced transcripts from this locus (Supplementary Figure 2). By combining RIP-seq enrichment with Nanopore results, we confirmed that EZH2 binds multiple intronic rather than exonic RNAs from this region (Figure 2A). Digital droplet PCR of K562 nuclear and cytoplasmic RNA further revealed that the EZH2-bound fragments of LEVER are nuclear-localized (Figure 2C).

### LEVER RNA sequesters PRC2 and inhibits H3K27 methylation at the LEVER locus

Given that loss of nascent transcripts globally increased PRC2-chromatin interactions and PRC2-related histone methylation levels (Figure 1E, 1F), we wondered if loss of specific transcripts would result in a similar observation. We thereby knocked out LEVER using CRISPR-Cas9 in K562 cells. To completely disrupt LEVER RNA generation while preserving corresponding gene loci for following epigenetic studies, we designed single guide RNAs targeting the LEVER promoter region in order to generate small deletions that disrupted RNA Pol II initiation at the LEVER locus (Supplementary Figure 3A). We obtained a LEVER-silenced clone with a homozygous deletion (270bp) spanning the TSS site (Figure 3A, Supplementary Figure 3B). Loss of LEVER transcripts increased EZH2 occupancy at the locus (Figure 3B), which was accompanied by markedly elevated H3K27me3 levels at the same loci (Figure 3C). In contrast, no noticeable increase in either EZH2 or H3K27me3 was detected at MYT1 (serving as a positive control for EZH2 and H3K27me3 ChIP) or CCDC26 (negative control) regions, suggesting the PRC2 related chromatin changes at the LEVER locus is site-specific upon LEVER RNA knock-out. To validate the above observations, we further constructed a doxycycline-inducible LEVER knock-down K562 line using the CRISPRi-dCas9 system. Of note, the dCas9 we used in our CRISPRi ^15^ experiments is not KRAB-fused ^16^, and thus the RNA knock-down effect is achieved by blockage of Pol II at the LEVER promoter without introducing any direct epigenetic modifications (Supplementary Figure 3A). Consistently, inducible knock-down of LEVER (Figure 3D) remodeled the PRC2-related epigenetic features along the LEVER locus but not nearby regions. Histone ChIP-seq showed increased H3K27me2/me3 and decreased H3K27ac at the LEVER locus (Figure 3E), suggesting locally enhanced PRC2 activity and reduced chromatin accessibility. These ChIP-seq results were further validated at selected regions by qPCR (Supplementary Figure 3C, 3D, 3E). Collectively, the depletion of LEVER transcripts specifically increased PRC2 occupancy and its progressive H3K27 methylation products on the local chromatin. This gene-specific observation reveals that LEVER RNA regulates the homeostasis of PRC2 related histone modifications *in cis* and is consistent with the above observations from global RNA depletion (Figure 1D, 1E, 1F).

**Figure 3.**
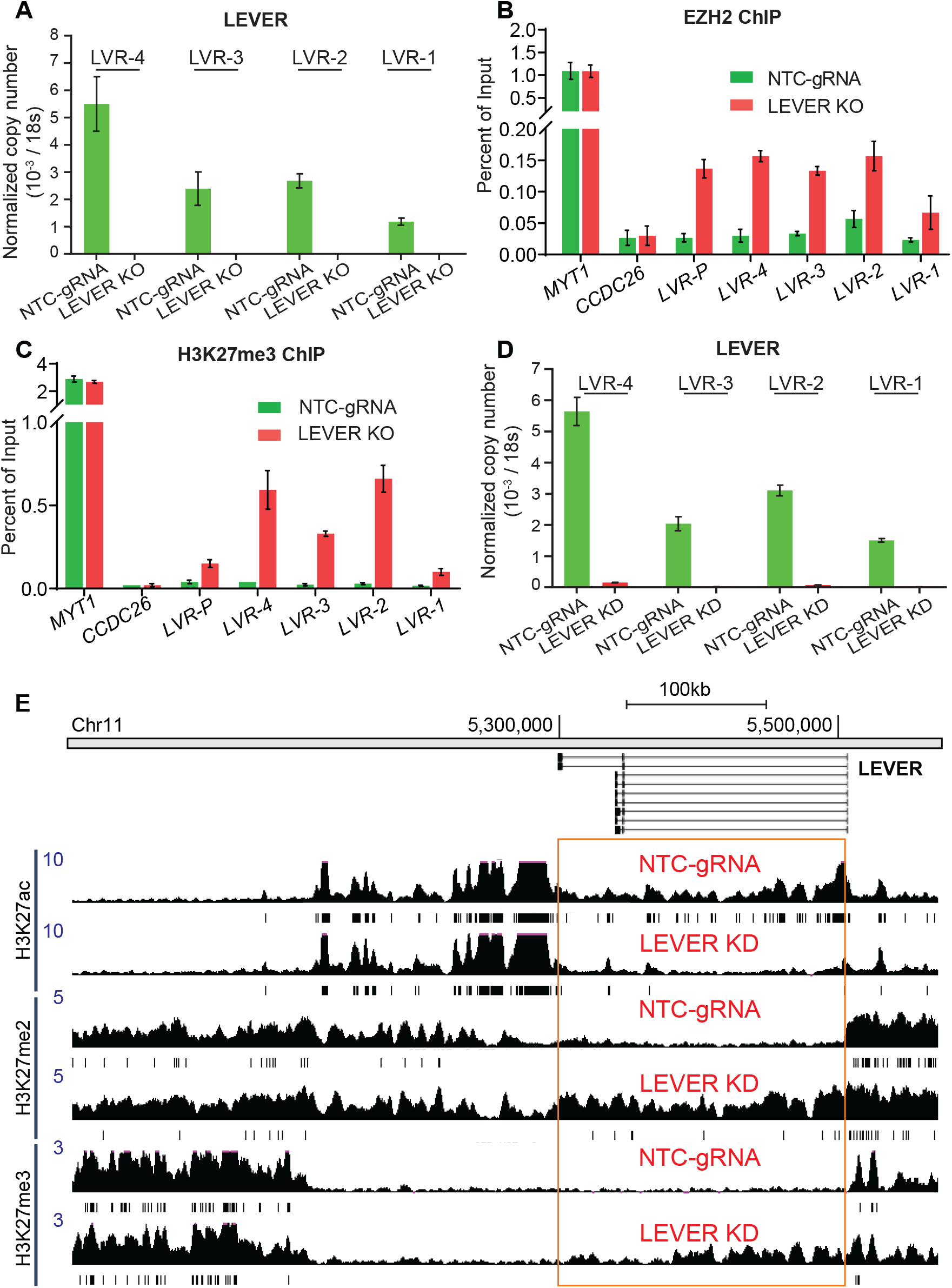
LEVER RNA sequesters EZH2 and blocks histone H3K27 methylation at the LEVER locus *in cis.* (A) LEVER RNA expression measured by primers detecting four selected regions (marked in Figure 2A) in non-targeting control (NTC-gRNA) or LEVER knock-out (LEVER KO) K562 cells. LEVER RNA levels were below the level of detection following LEVER knock-out. (B) (C) EZH2 (B) or H3K27me3 (C) chromatin occupation at selected regions within the LEVER locus. MYT1 and CCDC26 serve as ChIP positive and negative controls, respectively. LVR-P primer pair is designed at the LEVER promoter region (Supplementary Figure 3A). (D) LEVER RNA expression in doxycycline (dox) inducible non-targeting control (NTC-gRNA) or LEVER knock-down (LEVER KD) K562 cells treated with dox for 28 days quantified by RT-qPCR. (E) H3K27ac, H3K27me2, and H3K27me3 ChIP-seq at the LEVER locus in dox-inducible non-targeting control (NTC-gRNA) or LEVER knock-down (LEVER KD) K562 cells treated with dox for 28 days.

### LEVER RNA negatively regulates ε-globin

LEVER-PRC2 interaction and its epigenetic effects *in cis* may regulate local gene transcription. Due to the genomic proximity of LEVER to the β-globin cluster, we examined the regulation of expression of ε-globin, the β-globin member adjacent to LEVER (Figure 4A), by LEVER RNA. Knock-out and knock-down of LEVER RNA upregulated ε-globin mRNA levels (Figure 4B), and this increase was further observed at the protein level (Figure 4C). In contrast, γ-globin mRNA level was not affected by the loss of LEVER (Supplementary Figure 4A). Of note, although lncRNAs have been reported to affect mRNA stability ^17^, the increase of ε-globin mRNA abundance here was achieved by transcriptional activation, as reflected by increased RNA Pol II occupancy at ε-globin promoter upon LEVER knock-down (Figure 4D).

**Figure 4.**
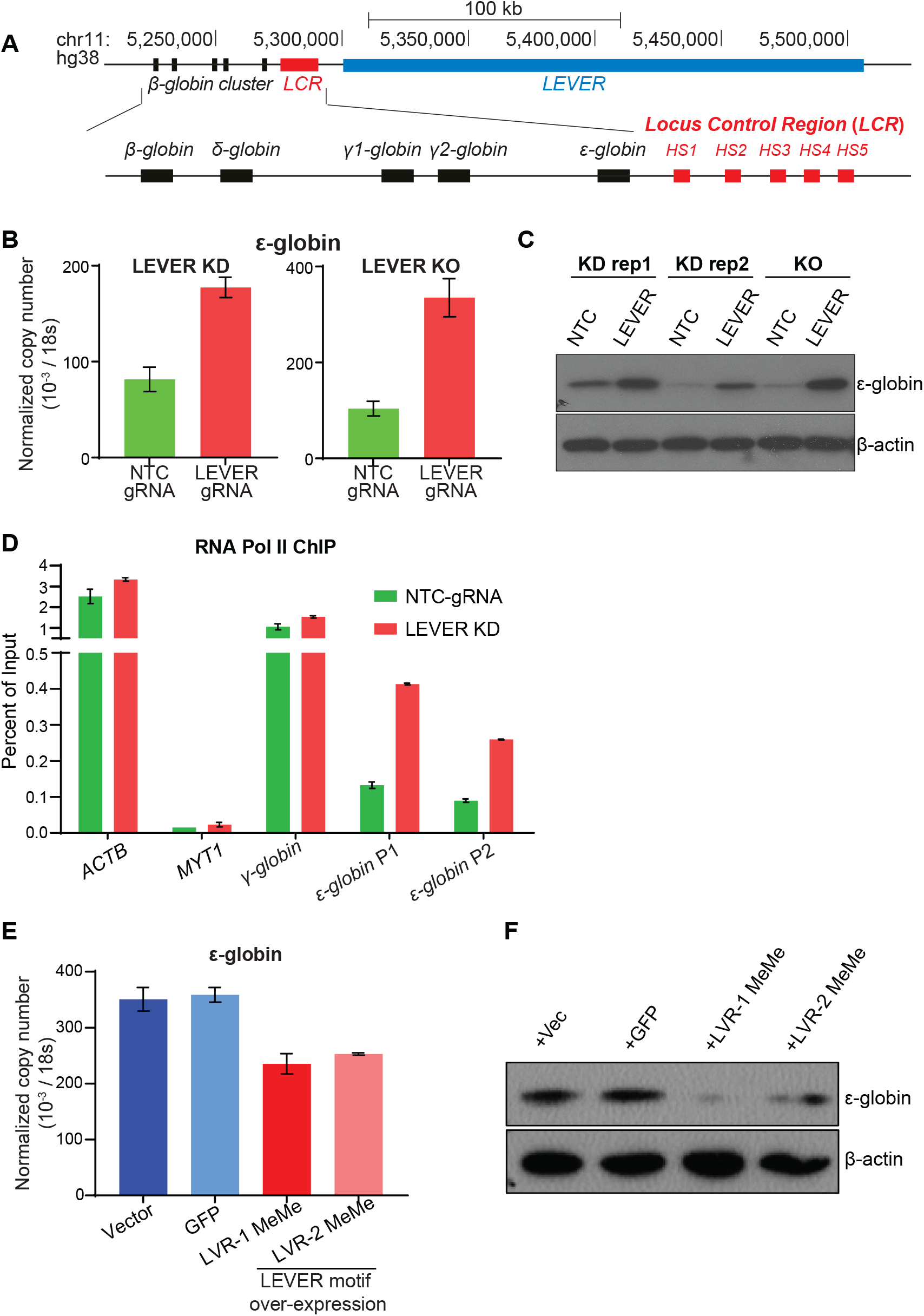
LEVER RNA negatively regulates ε-globin. (A) Chromosome map of the β-globin cluster and LEVER locus. Transcription of LEVER (in blue) and the globin genes (in black) proceeds in a right to left direction. HS1 through HS5 (in red) correspond to the hypersensitive regions in the LCR. (B) ε-globin expression after inducible LEVER knock-down (KD) or LEVER knock-out (KO) in K562 cells measured by RT-qPCR. (C) ε-globin protein abundance in LEVER KD or KO K562 cells measured by western blot. β-actin serves as the loading control. (D) ChIP-qPCR of RNA Pol II at ε-globin and γ-globin promoter in non-targeting control (NTC-gRNA) and LEVER knock-down (LEVER KD) cells. ACTB and MYT1 serve as Pol II ChIP positive and negative controls, respectively. Pol II occupation at ε-globin promoter is measured by two primer sets (ε-globin P1 and ε-globin P2). (E, F) ε-globin expression in LEVER KO K562 cells rescued by LVR-1 MeMe and LVR-2 MeMe. LEVER KO K562 cells were transduced with empty vector, GFP, LVR-1 MeMe, or LVR-2 MeMe expressing lentivirus and analysed for ε-globin expression using RT-qPCR (E) and western blot (F). LVR-1 and LVR-2 MeMe represent RNA fragments encompassing MeMe enriched EZH2-interacting RNA motif (Figure 1C) within the LVR-1 and LVR-2 regions.

To further validate the regulatory effect of LEVER RNA on ε-globin expression, we over-expressed two RNA fragments encompassing the consensus EZH2-bound guanidine-rich motif (Figure 2A) within LVR-1 and LVR-2 regions (termed LVR-1 MeMe and LVR-2 MeMe respectively) independently in LEVER KO cells. Restored expression of EZH2-interacting LEVER fragments (Supplementary Figure 4B) mitigated ε-globin activation at mRNA (Figure 4E) and protein levels (Figure 4F), showing an inverse correlation and supports the negative regulatory effect of LEVER RNA on ε-globin.

Moreover, we observed that the de-repression of LEVER RNA upon doxycycline withdrawal was associated with a reversible decrease in ε-globin RNA (Figure 5A; Supplementary Figure 4C). In sum, these dynamic studies clearly support a negative regulatory effect of LEVER RNA on ε-globin expression.

**Figure 5.**
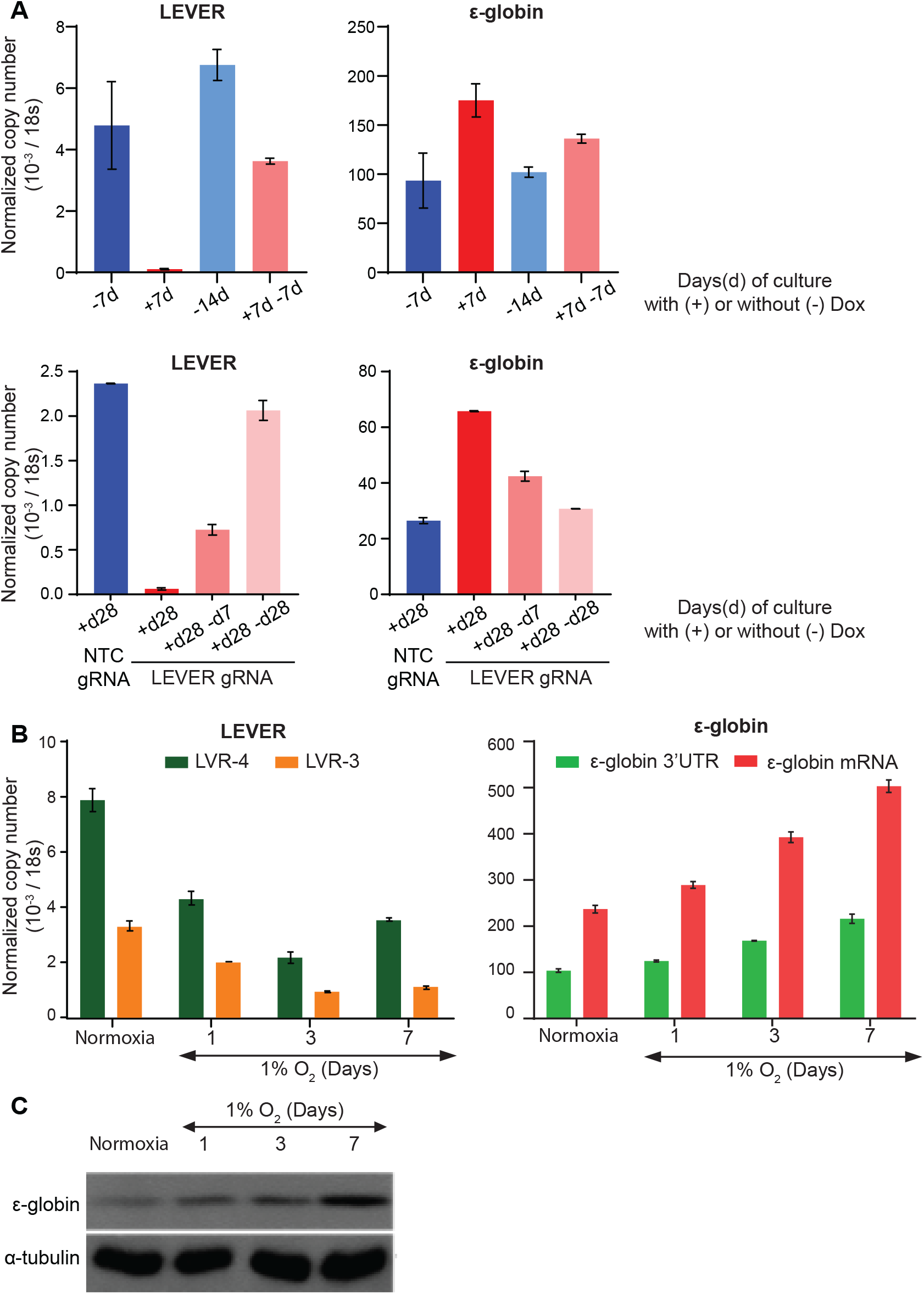
Dynamic LEVER regulation validates inverse correlation between LEVER and ε-globin expression. (A) LEVER and ε-globin expression in inducible LEVER KD K562 cells with (+) or without (-) doxycycline treatment for the indicated duration. LEVER expression is quantified at the LVR-4 region. (B) Time course RT-qPCR analysis of LEVER RNA and ε-globin in K562 cells during 7-day hypoxic culture under 1% oxygen. (C) ε-globin protein abundance in K562 cells cultured under 1% oxygen. α-tubulin serves as the loading control.

### Hypoxia regulates LEVER and ε-globin

The embryonic form of β-globin, ε-globin, is expressed almost exclusively in the early human embryo in which the environment is hypoxic ^18^. To study whether LEVER can be a *bona fide* regulator of ε-globin, we examined the expression homeostasis between these two under environmental perturbations, e.g., oxygen levels. Culturing K562 cells in hypoxic conditions (1% oxygen) repressed LEVER while enhanced ε-globin expression over time (Figure 5B, 5C), which reproduced the inverse-correlation between LEVER and ε-globin expression. Of note, LEVER RNA level was more drastically changed in terms of fold difference than ε-globin on the first day of hypoxic culture, suggesting LEVER as an upstream regulator of ε-globin.

### The LEVER locus regulates ε-globin through long-range chromatin interactions

Although we have demonstrated that LEVER RNA inhibits EZH2 function at its own locus *in cis*, LEVER RNA may also act directly through PRC2 to repress the ε-globin locus *in trans.* We thereby examined the H3K27me3 status at the ε-globin locus. We did not detect noticeable H3K27me3 coverage at the ε-globin locus at baseline. This observation suggests that ε-globin is not a direct repressive target of PRC2 mediated by H3K27 methylation in K562 cells.

To understand the role of EZH2 and LEVER in ε-globin transcriptional regulation, we knocked out EZH2 in K562 cells. Interestingly, ε-globin expression was reduced in EZH2 null cells in which global H3K27me3 was not detectable by western blot (Figure 6A, 6B). Re-expression of wild type EZH2 and its activating mutant (EZH2^Y641F^) ^19, 20^ in the knock-out cells restored ε-globin expression, while the truncated EZH2 mutant without methyltransferase motif (EZH2^ΔSET^) failed to do so (Figure 6A, 6B). In addition, the EZH2 enzymatic inhibitor GSK343 curtailed ε-globin up-regulation in LEVER knock-out cells (Supplementary Figure 5). In summary, this data suggests that this ε-globin activating effect requires EZH2 methyltransferase activity. Moreover, it appears that EZH2 function affected H3K27 methylation within the LEVER locus but not at the ε-globin gene. These results collectively suggest that EZH2 and LEVER regulate ε-globin in K562 cells.

**Figure 6.**
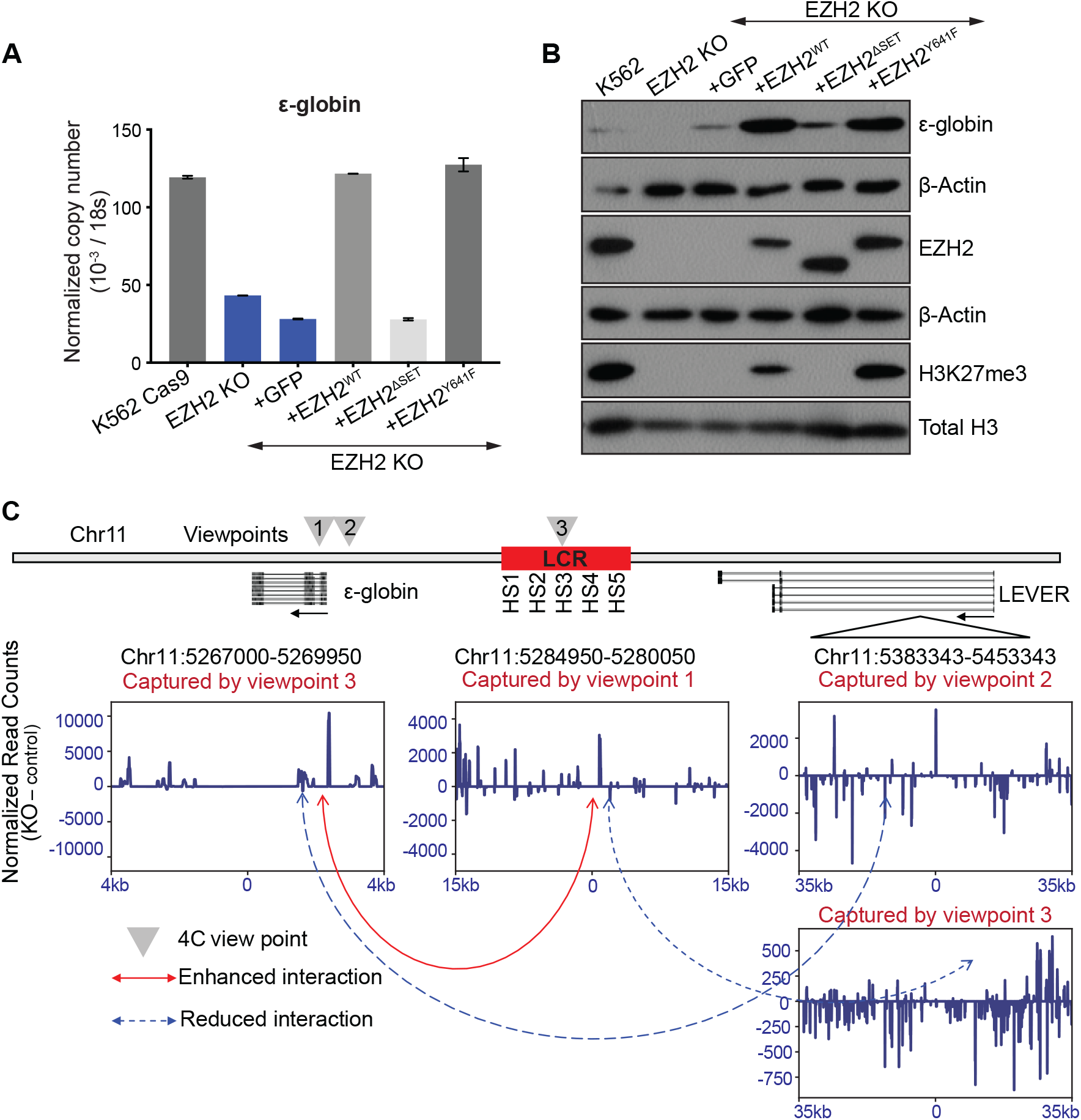
The LEVER locus regulates ε-globin through long-range chromatin interactions. (A) RT-qPCR analysis of ε-globin in K562 Cas9-expressing parental cells, EZH2 knock-out (EZH2-KO) cells, and EZH2-KO cells rescued with GFP, EZH2, EZH2^ΔSET^ and EZH2^Y641F^ EZH2^ΔSET^, truncated EZH2 without methyltransferase enzymatic domain; EZH2^Y641F^, EZH2 gain of methyltransferase enzymatic function mutation. The EZH2-KO cells were transduced with lentivirus to stably express these constructs. (B) Western blot analysis of ε-globin, EZH2, and global H3K27me3 in K562 wild type, EZH2 KO, and rescue cells as in (A). β-actin and total histone H3 were used as loading controls. (C) 4C analysis comparing local chromatin interactions near the β-globin cluster in LEVER KO and NTC cells shows increased ε-globin-LCR interaction upon LEVER knock-out. We positioned two viewpoints at the ε-globin promoter and two viewpoints at HS3 and HS4 regions within the β-globin locus control region (LCR). The viewpoints at ε-globin promoters and HS3 are shown here and indicated as viewpoints 1-3, respectively. Captured genomic fragments displayed in the windows correspond to the genomic coordinates annotated above each window. Differentially interacted regions are shown in subtracted reads by comparing LEVER KO and NTC cells, thereby positive or negative values represent increased (solid red line) or decreased (dotted blue lines) interactions upon LEVER KO, respectively.

The β-globin cluster is well known to be regulated by long-range chromatin interactions. Therefore, we asked whether the LEVER locus itself may work as a negative regulatory element of the ε-globin gene through long-range chromatin interactions. To test this hypothesis, we performed 4C-seq analysis before and after LEVER promoter knock-out to compare local chromatin interaction changes upon LEVER RNA loss. We positioned two viewpoints at the ε-globin promoter, and another two viewpoints at HS3 and HS4 regions within the β-globin Locus Control Region (LCR), which are well-established activating elements of the β-globin cluster ^21^ (Figure 6C). As expected, our 4C-seq analysis suggested that the ε-globin regulatory domains, including both the ε-globin promoter and the two LCR regions, interact weakly but broadly to LEVER chromatin in K562 cells. Loss of LEVER RNA resulted in the reduction of such interaction (Figure 6C, Supplementary Figure 6), presumably due to the compromised accessibility of LEVER locus caused by PRC2 methylation. In addition, the interaction between ε-globin promoter and LCR regions was significantly enhanced upon LEVER KO, as supported by the results from both viewpoints (Figure 6C, Supplementary Figure 6). Together with the increased Pol II occupancy at ε-globin promoter (Figure 4D), this observation suggests that the strengthened ε-globin promoter-LCR enhancer interaction could be the cause of ε-globin transcriptional activation upon LEVER RNA loss. Since the LEVER locus interacts with both the ε-globin promoter and the LCR, we propose that it may work as a competitor of the ε-globin promoter for LCR binding, thereby exerting its negative regulatory effect on ε-globin expression. Collectively, our 4C-seq results from different viewpoints support LEVER RNA as a negative regulator of ε-globin.

## Discussion

Following the initial discovery of the specific PRC2-lncRNA interaction and its supportive effect on PRC2 functions at PRC2 target genes ^2–4, 11^, more pieces of evidence revealed a promiscuous PRC2-nascent RNA interaction that inhibits local PRC2 function ^5–8,14^. While previous studies of the latter viewpoint were performed either *in vitro* or exclusively in mESC, our study in the human chronic myelogenous leukemia cell line K562 provides further evidence to support this model. In summary, we demonstrated that PRC2 manifests genome-wide interaction with nuclear-localized, non-polyA tailed, guanidine-rich intronic RNAs that are transcribed from active gene loci. These genes, as suggested by local H3K27me3 ChIP-seq profiling, tend to localize within densely H3K27 methylated genomic regions, but they are rather sparsely methylated within their gene body. These observations suggest that the PRC2-sequestering function of their nascent transcripts may play a role in antagonizing PRC2 repression *in cis* within PRC2-dense environment. Following the global analysis, we further identified and characterized the LEVER locus that generates PRC2-binding nascent transcripts near the β-globin cluster. Using both CRISPR knock-out and inducible CRISPRi knock-down systems, we showed that LEVER RNA prevents PRC2 from occupying the chromatin and thereby curtails EZH2 methyltransferase activity at the LEVER locus (Figure 7). These observations made at single gene locus further support the model generalized from global analysis that nascent transcripts deprive PRC2 from the chromatin and block PRC2 function locally.

**Figure 7.**
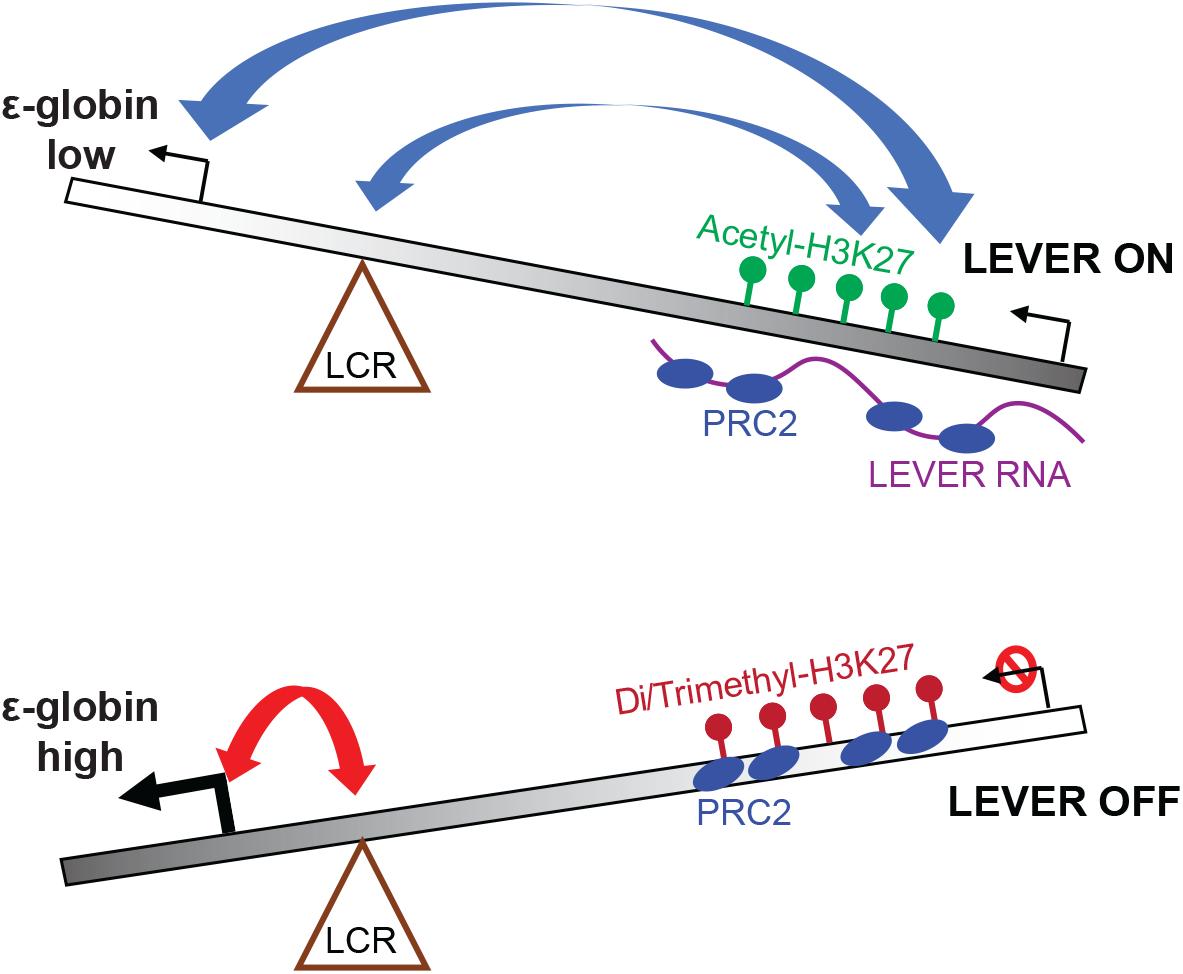
The proposed model of LEVER RNA regulating ε-globin transcription. Top diagram: LEVER RNA sequesters and antagonises PRC2 *in cis.* This maintains the accessibility of the LEVER locus and facilitates its chromatin interaction with the ε-globin promoter and LCR region, which negatively regulates ε-globin transcription. Bottom: When LEVER is silenced, the LCR can now interact with the ε-globin promoter to induce ε-globin transcription.

Our study further reveals a novel model of epigenetic regulation on ε-globin expression (Figure 7). Our results showed that in the presence of the LEVER RNA, the LEVER locus competes with the ε-globin promoter to bind with the LCR, a well-established enhancer region for the β-globin cluster ^21–22^, and thereby suppresses ε-globin transcription. In the absence of LEVER RNA, the released PRC2 occupies the LEVER chromatin, facilitates H3K27 methylation, and reduces the accessibility of the LEVER locus. This in turn permits the interaction between the LCR and the ε-globin promoter, which promotes the transcription of ε-globin. In sum, we demonstrated that the LEVER RNA can activate the LEVER locus that works as a negative regulatory element of ε-globin gene expression through chromatin looping.

Interestingly, we found that transcripts generated from similar genomic regions upstream of the β-globin cluster in the mouse genome were previously identified and annotated as polycomb-associated ncRNAs, similar to LEVER ^23^. But the two loci from the two species share poor sequence homology (data not shown). We wonder whether the regulatory effect of the corresponding LEVER loci and RNAs on β-globin cluster are functionally conserved in mouse although the loci are poorly conserved. Considering the promiscuous RNA binding pattern of PRC2, some PRC2-binding non-coding RNAs may have evolved to become opportunistic regulatory elements that are epigenetically functional to neighboring genes, and thereby conserved their relative position to neighboring protein-coding genes without the need to preserve their sequences in different species. The syntenic transcription of lncRNA has been proposed together with conserved sequence, structure, and function as the four dimensions of lncRNA conservation ^24^ Our model in PRC2-nascent RNA interaction and its regulatory function on nearby genes may explain such synteny conservation observed in some lncRNAs.

In addition, our identification of the LEVER locus as a negative regulator of ε-globin shows potential clinical significance. β-hemoglobinopathies, including sickle cell disease (SCD) and β-thalassemia, are caused by genetic mutations in the adult-form β-globin gene. They are among the most common genetic diseases and influence tens of millions of patients worldwide ^25^. Reactivation of the fetal γ-globin gene has been demonstrated to ameliorate the severity of these diseases ^26^. Therefore, much effort has been made to understand the molecular basis of γ-globin regulation and to develop therapeutic strategies to reactivate γ-globin expression in β-hemoglobinopathy patients ^27–28^ However, the embryonic-form globin, ε-globin, is much less studied than its fetal counterpart ^29^, even though limited studies indeed suggested the therapeutic potential of ε-globin reactivation in the SCD mouse model, similar to γ-globin ^30, 31^. In this work, we provided an in-depth understanding of the regulation of ε-globin by its upstream LEVER loci through PRC2. More importantly, we have achieved multi-fold ε-globin upregulation through knocking-out and knocking-down the non-coding transcript from LEVER. This moderate enhancement of ε-globin expression, however, may ameliorate red blood cell sickling and thereby mitigate SCD symptoms as suggested by previous studies ^30, 31^. Taken together, our findings suggest a novel therapeutic strategy of using epigenetic modulators, RNAi, or even gene editing to reactivate ε-globin for the treatment of β-hemoglobinopathy patients.

---

**Supplementary Figure 1.**
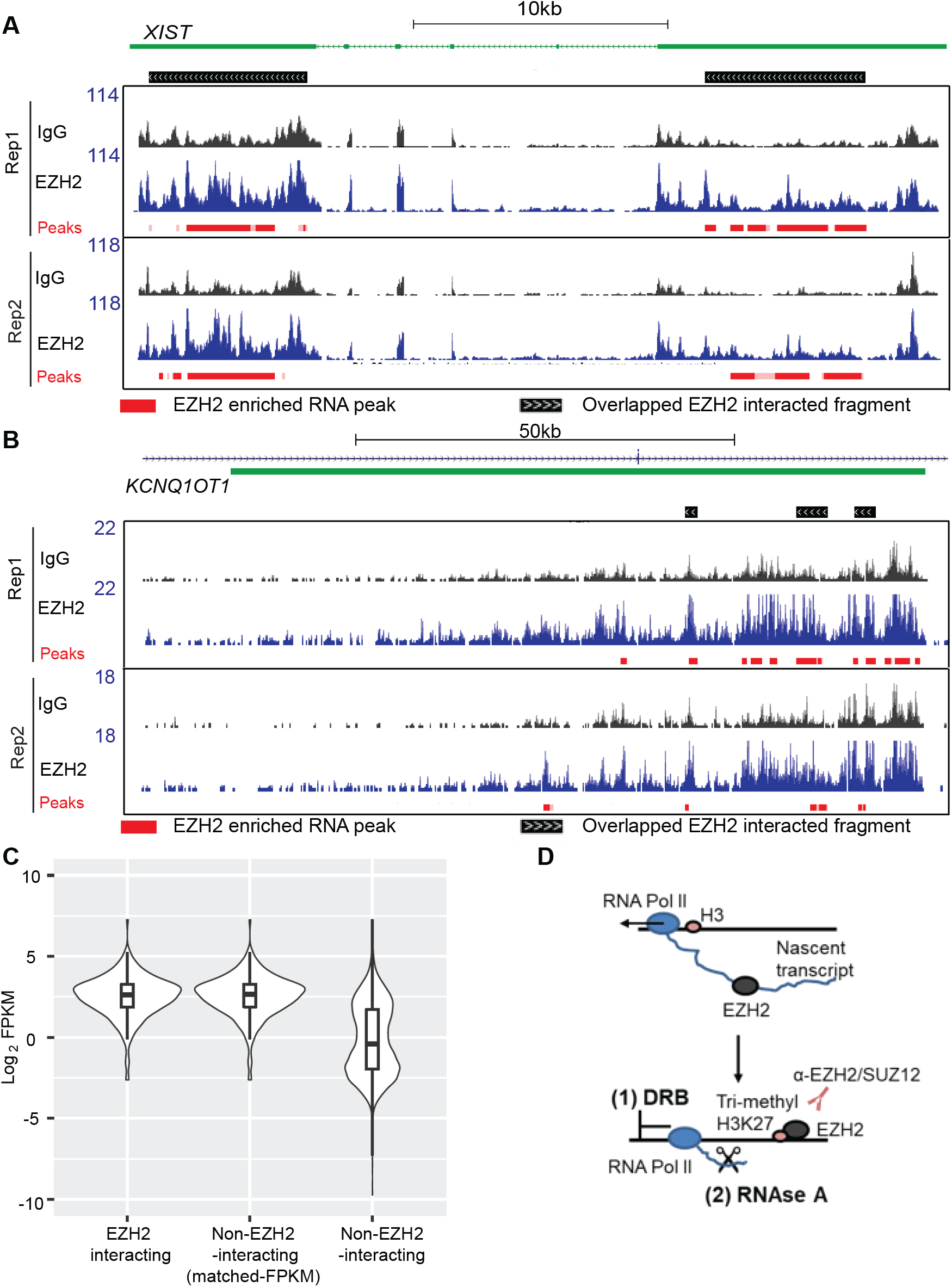
(A) Genome tracks of enriched EZH2 RIP peaks of XIST RNA with reference to Genecode v29 in K562 cells. (B) Genome tracks of enriched EZH2 RIP peaks of KCNQ1OT1 RNA with reference to Genecode v29 in K562 cells. (C) Log2 converted FPKM values of EZH2-interacting RNAs, a FPKM-matched subset of non-EZH2-interacting RNAs, and the total non-EZH2 interacting RNAs. (D) Design of experiments to demonstrate dependence on transcription and/or RNA (Figure 1E and 1F). K562 cells were treated with (1) 100μM DRB or vehicle (DMSO); or (2) 1 mg/ml RNase A or vehicle. Immunoprecipitation of EZH2 or SUZ12 was performed after treatment to analyse the binding of PRC2 to histone H3.

**Supplementary Figure 2.**
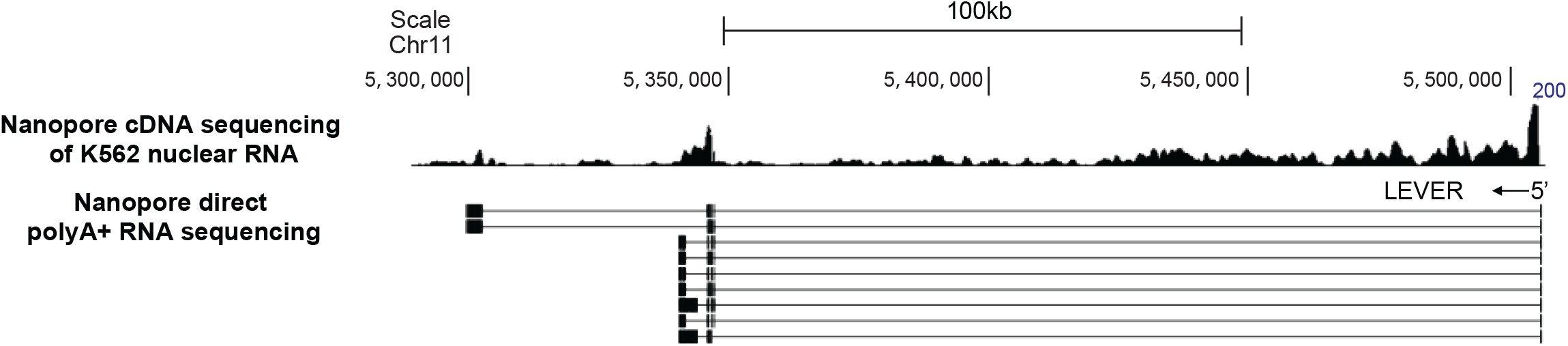
Mapping of LEVER RNA to the human genome with Nanopore cDNA sequencing of K562 nuclear RNA or direct-RNA sequencing of K562 total polyA+ RNA.

**Supplementary Figure 3.**
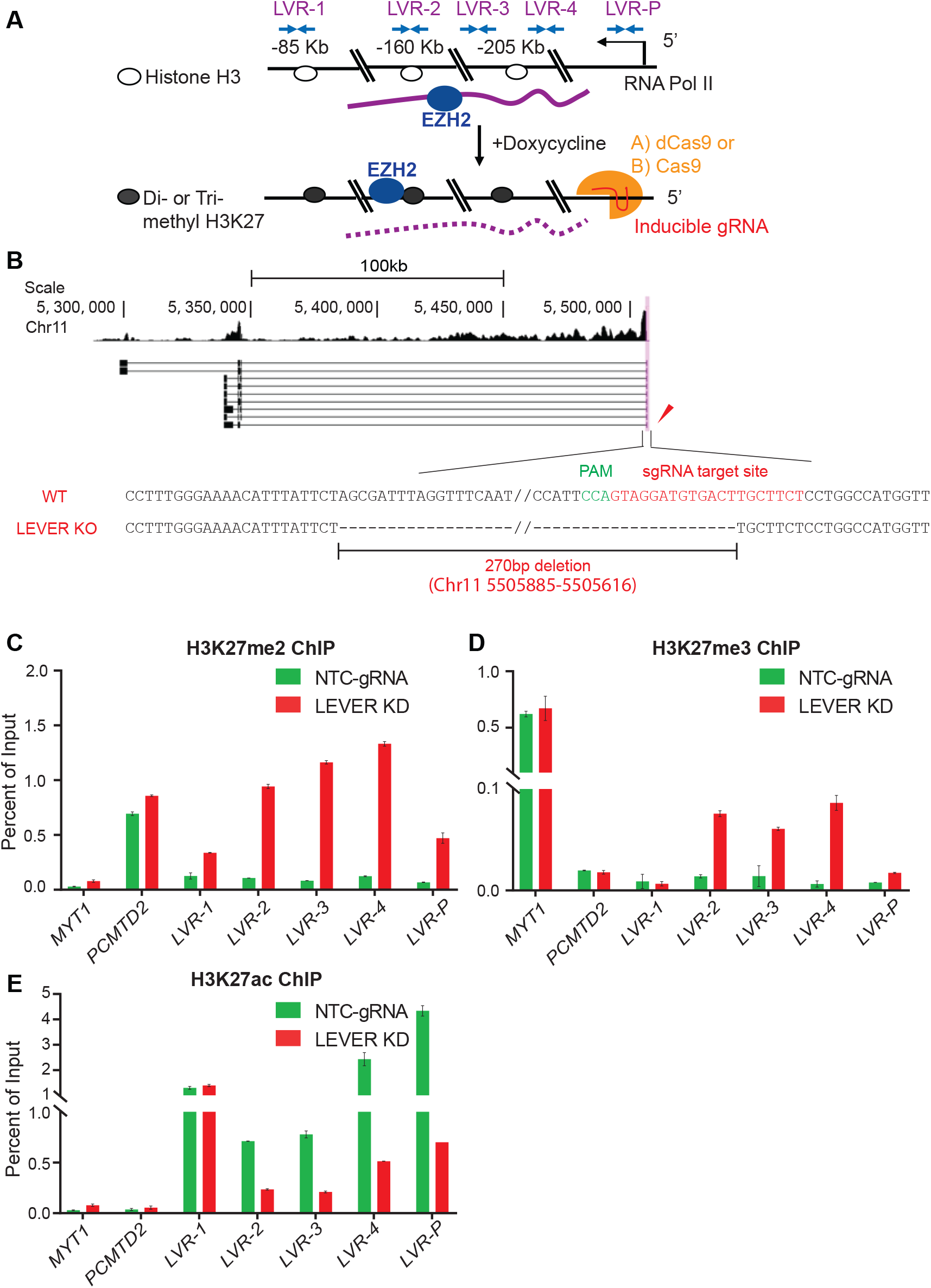
(A) Illustration of LEVER knockout (KO) or doxycycline-inducible knock down (KD) using the CRISPR-Cas9 or CRISPRi-dCas9 system. The guide RNA was designed to target at the LEVER TSS region. (B) CRISPR-Cas9 generated LEVER KO cells with homozygous deletion spanning the LEVER promoter. (C,D,E) ChIP-qPCR analysis of (C) H3K27me2, (C) H3K27me3, and (D) H3K27ac along the LEVER locus in doxycycline treated LEVER KD cells or NTC cells after 28 days of doxycycline. MYT1 serves as a ChIP positive control and PCMTD2 serves as a ChIP negative control.

**Supplementary Figure 4.**
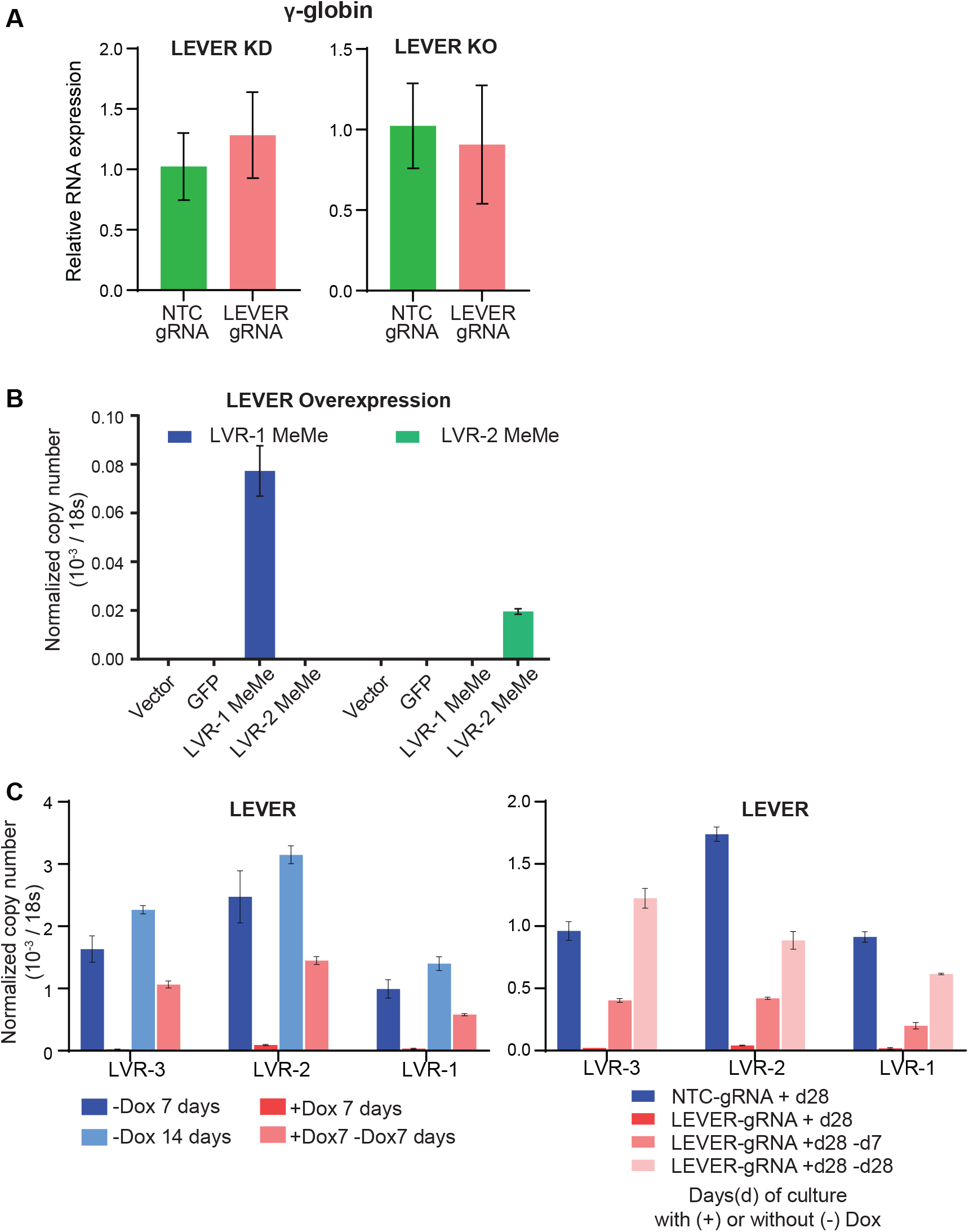
(A) RT-qPCR analysis of γ-globin in LEVER knock-down and knock-out cells. Notably, the β- and δ-globin genes are not stably expressed in K562 wild-type or LEVER depleted cells. (B) RT-qPCR analysis of LEVER-MeMe fragments in LEVER-KO+Vector control, KO+GFP control, KO+LVR-1 MeMe, and KO+LVR-2 MeMe. (C) RT-qPCR of LEVER expression at LVR-1, LVR-2, and LVR-3 regions in corresponding cells and culturing conditions for Figure 5A and 5B, respectively.

**Supplementary Figure 5.**
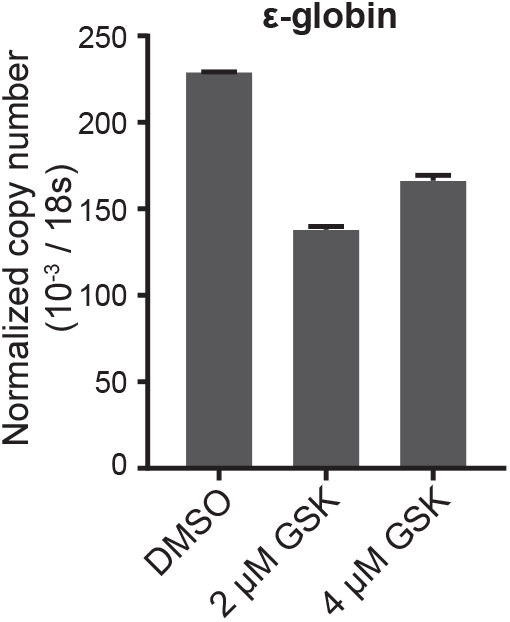
Quantitative RT-PCR of ε-globin in LEVER knock-out cells treated with DMSO, 2 μM GSK343, or 4 μM GSK343.

**Supplementary Figure 6.**
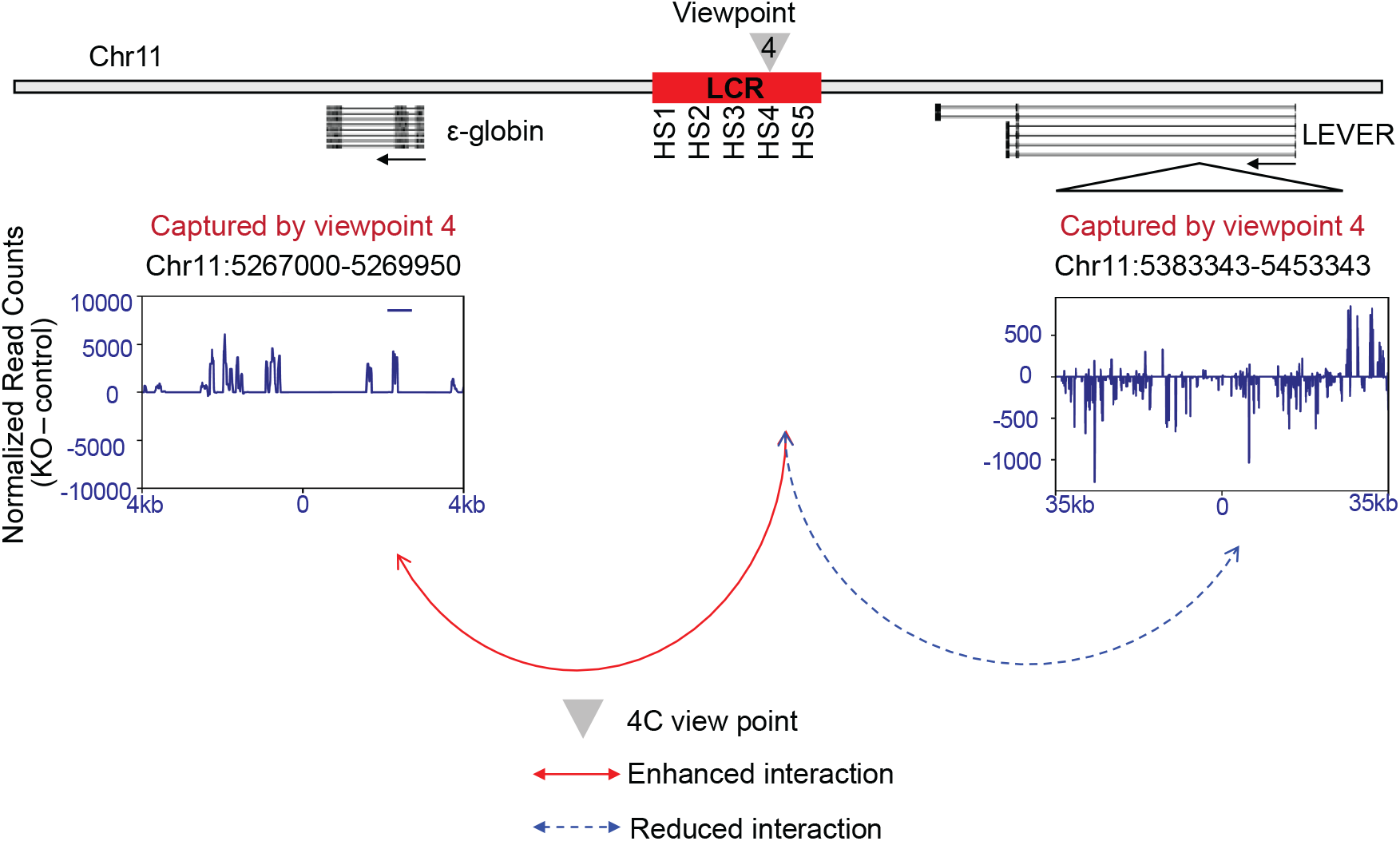
4C analysis of NTC and LEVER KO K562 cells at viewpoint 4 as described in Figure 6C.

## Materials and Methods

### Cell culture

K562 and HEK293T cell lines were obtained from the American Type Culture Collection (ATCC) and were used in experiments at their early passages. K562 cells were cultured in RPMI (Biowest) supplemented with 10% fetal bovine serum (FBS) (Biowest). HEK293T cells were cultured in DMEM supplemented with 4500 mg/L glucose (Biowest), 2mM L-glutamine (Thermo Scientific), 1X MEM non-essential amino acids (Thermo Scientific), 1mM sodium pyruvate, and 10% fetal bovine serum (FBS) (Biowest). All cell lines were tested for mycoplasma every six months.

### Lentivirus production

pCMV-dR8.91, pCMV-VSV-G, and lentivectors were transfected into 10 million 293T cells using Lipofectamine 2000 following the manufacturer’s instructions. Supernatants were harvested at 48 and 72 h after transfection. Viruses were concentrated 100–300 times through centrifugation after filtering through 0.45 μM syringe filters and stored at −80°C. The titers of viruses were determined on HeLa cells using flow cytometry. The plasmids used in this study are listed in Supplementary Table S2.

### CRISPR/Cas9 and CRISPRi/dCas9

K562 cells were transduced with the FUCas9Cherry or FUdCas9Cherry lentiviruses at 0.5 MOI with 5μg/mL polybrene. The transduced cells were sorted for mCherry positive as K562-Cas9/dCas9 stable lines. FgH1tUTG-sgRNA viruses were further transduced into these K562-Cas9/dCas9 stable lines and sorted for GFP positives. To generate EZH2 KO, in vitro synthesized sgRNA (Synthego) was transiently transfected into K562-Cas9 cells with DharmaFECT 1 (DHARMACON) following the manufacturer’s instructions. Of note, the DNA editing efficiency of the sgRNAs at their specific targeting regions was determined in K562-Cas9-sgRNA cells by the T7 endonuclease I (NEB) assay (as described in the manufacturer’s protocol) after sgRNA induction by 1μg/mL doxycycline for 3 days. EZH2-sgRNA transfected (48 hours) K562-Cas9 cells or Induced K562-Cas9-sgRNA cells were seeded into single-cell clones in 96-well plates. Successful KO clones screened by RT-qPCR (for LEVER KO) or western blot (for EZH2 KO) were expanded and analyzed by genotyping. The list of sgRNAs used can be found in Table S2.

### Ectopic EZH2 or LEVER expression

For the ectopic expression of EZH2 and its mutants, N-terminal Flag-tagged EZH2 (WT) was cloned from K562 cDNA into a modified pLX304 plasmid in which the CMV promoter was replaced with the CAG promoter. The EZH2^ΔSET^ and EZH2^Y641F^ mutants were derived from EZH2 WT and cloned into the same vector. For the ectopic expression of LEVER fragments, LVR-1 or LVR-2 were amplified from K562 genomic DNA and cloned into a modified pLX304CAG plasmid, in which the CAG promoter and the hPGK promoter were inverted, and a BGH stop sequence was added to ensure that only the designated LEVER RNA fragments are being expressed. Lentivirus was produced as stated above and transduced into corresponding knock-out K562 cells and selected with blasticidin.

### Western blot

Protein lysates were prepared in RIPA buffer (50mm Tris-Cl pH 7.5, 150 mM NaCl, 1 mM EDTA, 1% NP-40, 0.5% sodium deoxycholate (NaDOC), 0.1% SDS), and the lysates were mildly sonicated into clear supernatants before subjected to protein quantification by the Pierce BCA assay (Thermo Scientific). 10-20 μg of protein lysate was loaded into each well of SDS-PAGE mini gel. The electrophoresis was performed in Tris-glycine buffer supplemented with 0.1% SDS. Proteins were then transferred onto nitrocellulose membranes (Bio-rad) in Tris-glycine-methanol transfer buffer. The membrane was blocked by 5% skim milk/TBST and incubated with primary antibodies followed by HRP-conjugated secondary antibodies diluted in 5% skim milk/TBST. Chemiluminescence was detected by exposing X-ray film to the blot pre-incubated with the Luminata Crescendo Western HRP substrate (Millipore). The list of antibodies used can be found in Supplementary Table S1.

### Quantitative PCR (qPCR)

1 μg of RNA was reverse transcribed using Qscript cDNA Supermix (QuantaBio). qPCR on cDNA or ChIP DNA was performed with GoTaq qPCR Master Mix (Promega) under the manufacturer’s instruction on a QS5 system (Thermo Scientific). The copy numbers were measured by predetermined standard curves. The relative expression or enrichment to control samples were calculated based on the ΔΔCt method. Quantitative digital droplet PCR was performed on a QX200 Droplet Digital PCR (ddPCR) System (Bio-Rad) with EvaGreen Digital PCR Supermix (Bio-Rad) under the manufacturer’s instructions. Primers used in this study are listed in Supplementary Table S2.

### RNA fractionation and sequencing

Nuclear RNA purification was performed as described previously ^1^. Briefly, 12 million cells were harvested and washed first with 20mL of cold PBS supplemented with 5mM ribonucleoside Vanadyl complex (NEB) and 1mM phenylmethylsulphonyl fluoride (PMSF), and then with 20mL of cold PBS supplemented with 1mM PMSF. The cell pellet was lysed in 1.7 mL of cold hypotonic buffer (10mM HEPES pH7.6, 25mM KCl, 0.15mM spermine, 0.5mM spermidine, 1mM EDTA, 2mM sodium butyrate, 1.25M sucrose, 10% glycerol, 5mg/mL BSA, 0.5% NP-40) supplemented with 1mM PMSF, EDTA-free protease inhibitor cocktail (Roche) and 20U/mL SUPERase-In (Invitrogen). The cells were homogenized with a 15 mL Dounce homogenizer (by 10 strokes with Pestle A and 45 strokes with Pestle B). The homogenized cells were diluted in 7.8 mL of NP-40 free hypotonic buffer containing 2M sucrose. 3.2 mL of cushion buffer (BSA and NP-40 free hypotonic buffer containing 2M sucrose) was added to the bottom of a polypropylene tube (Beckman Coulter, 331372), and the cell lysate was overlaid on the cushion buffer without disrupting the bottom phase. Nuclei were harvested after ultra-centrifugation at 100,000 x g for one hour at 4°C with a SW41Ti rotor and Beckman Coulter TH641 ultra-centrifuge. The supernatant was removed, and the nuclear RNA was extracted by Trizol according to the manufacturer’s protocol. Nuclear polyA+ RNA was extracted with a NEBNext Poly(A) mRNA Magnetic Isolation Module according to the manufacturer’s protocol, and the nuclear polyA-fraction was extracted from the unbound supernatant by ethanol precipitation. Cytoplasmic RNA was isolated with a Qiagen RNeasy kit according to the manufacturer’s protocol. The fractionated RNA was subjected to DNase I digestion and then re-purified using Trizol. The RNA was quantified with the Qubit RNA HS assay kit (Thermo Scientific), and rRNA was removed by the Ribo-Zero Gold rRNA Removal Kit (Illumina). The stranded RNA-seq libraries were constructed with the Script-seq V2 RNA-seq library preparation kit (Illumina). The 250–500 bp size-selected libraries were sequenced on the Illumina MiniSeq platform with MiniSeq High Output 2X150 paired-end kit.

### DRB / RNase A treatment and immunoprecipitation

For DRB treatment, 12 million K562 cells were treated with 100 μM DRB for 24 hours before subjected to nuclei extraction. For RNase A treatment, 10 million K562 cells were first harvested and washed with 1X PBS, and then treated with 1 mg/mL RNase A in 0.05% Tween-20 for 10 mins at RT before subjected to nuclei extraction. The DRB/RNase A treated cells were lysed on ice for 5 min with cell lysis buffer (20 mM Tris pH 7.5, 10 mM KCl, 1.5 mM MgCl_2_, 0.1% Triton X-100) supplemented with EDTA-free protease inhibitor cocktail (Roche), phosphatase inhibitor (Roche), 1 mM DTT, and 1 mM PMSF and then centrifuged at 800 RPM for 10min at 4°C. The nuclei pellets were washed once with the cell lysis buffer without Triton X-100 and harvested by centrifugation at 1300 x g for 4 min at 4°C. Nuclei were resuspended in resuspension buffer (50 mM Tris pH 7.5, 150 mM NaCl, 2 mM EDTA) supplemented with EDTA-free protease inhibitor cocktail (Roche), phosphatase inhibitor (Roche), 80 U/mL RNaseOUT (ThermoScietific), and 1 mM PMSF. The nuclei were sonicated using EpiShear Probe Sonicator (Active Motif) for 15 seconds at 30% amplitude. 500 μl of the lysate was incubated with EZH2 antibody (1:250), SUZ12 antibody (2 μg), or rabbit IgG isotype (2 μg) in the cold room for 12 hours. The lysate was incubated with 50 μl pre-blocked dynabeads protein A (Invitrogen) for 2 hours at 4 °C. The beads were washed with washing buffer (150mM NaCl, 10mM Tris-HCl, pH7.4, 1mM EDTA, 1mM EGTA, pH8.0, 1% Triton X-100, 0.5% NP-40, 1U/mL SUPERase-In) for three times before eluted with 30 μl of RIPA buffer and 15 μl of 6X Laemmli buffer at 95°C for 10 mins. The eluted complex was analyzed with western blot.

### RNA immunoprecipitation (RIP)

RIP was performed as described previously ^1^ with the following modifications. (A) SUPERase-In was used in substitution of ribonucleoside Vanadyl complex from the cell lysis step. (B) Fixed K562 cells were lysed with Dounce homogenizer by 10 strokes using Pestle A and 60 strokes using Pestle B. (C) The nuclei were sonicated with Bioruptor (Diagenode) for 8 cycles (30s on, 30s off, high power). (D) Nuclear lysates were pre-cleared with Dynabeads protein G twice at 4°C. (E) 5 μg of EZH2 or IgG isotype control antibodies were used in each RIP experiment. RIP experiments were performed in three biological replicates, and two of these replicates were independently sequenced. For sequencing, the immunoprecipitated RNA was quantified with the Qubit RNA HS assay kit (Thermo Scientific) before rRNA Removal with Ribo-Zero Gold rRNA Removal Kit (Illumina). 15ng of the rRNA-depleted RNA was used for sequencing library construction. The stranded RIP-seq libraries were constructed with a Script-seq V2 RNA-seq library preparation kit (Illumina). The libraries were subjected to size selection (250–500 bp) on a 4–20% TBE PAGE gel (Thermo Scientific). The recovered libraries were sequenced on the Illumina MiniSeq platform using the MiniSeq High Output kit with 2×150 paired-end.

### Chromatin immunoprecipitation (ChIP)

ChIP was performed as described previously ^2^ with minimal modifications. Briefly, K562 chromatin was sonicated with Bioruptor (Diagenode) for 15-25 cycles (30s on, 30s off, high power), and immunoprecipitation was performed with Dynabeads protein G. For ChIP-sequencing, the libraries were constructed using a Thru-PLEX DNA-seq 12S kit (TakaraBio). The libraries were subjected to size selection (250–500 bp) on a 4–20% TBE PAGE gel (Thermo Scientific). The recovered libraries were sequenced on the Illumina Nextseq platform using Nextseq 500 high-output kit with 2 × 76 paired-end. For H3K27me3, H3K27me2, or H3K27Ac ChIP-seq, one replicate was sequenced, and the results were validated by ChIP-qPCR with independent biological replicates.

### Oxford Nanopore sequencing

Fractionated nuclear RNA was used to construct a cDNA library using a 1D sequencing kit (SQK-LSK108) according to the manufacturer’s instructions. In brief, first-strand cDNA was generated from 20 ng of rRNA depleted nuclear RNA using SuperScript IV reverse transcriptase with 0.35 pmol/μl random hexamer at 52°C for 60 mins. Second strand cDNA was synthesized with the NEBNext second strand synthesis kit for 1 hour at 16°C. cDNA was purified with AMPure XP bead. Library was constructed with LongAmp Taq PCR for 18 cycles. The PCR product was purified with phenol: chloroform: isoamyl alcohol (25:24:1) and ethanol precipitated in the presence of glycogen and was finally dissolved in 10 mM Tris buffer (pH8). The cDNA library was sequenced on a Nanopore flow cell (FLO-MIN106). Enriched polyA+ RNA was used to construct the library for direct RNA sequencing (SQK-RNA001) according to the manufacturer’s instructions. In brief, 400 ng of total polyA+ RNA was used for reverse transcription adaptor (RTA) ligation before subjected to reverse transcription. RNA/DNA hybrids were purified with RNAClean XP beads and ligated with RNA adaptors as direct RNA sequencing library. The constructed library was sequenced on the Nanopore flow cell (FLO-MIN106).

### Circularized Chromosome Conformation Capture (4C)

4C-seq was performed as described previously ^3^ with modifications. K562 cells were fixed as ChIP protocol except that the cells were fixed in 2% formaldehyde at the concentration of 1 million cells per mL for 10min. Cells were lysed in complete lysis buffer (10 mM Tris-HCl, pH 8, 10 mM NaCl, 0.2% NP-40, and complete protease inhibitor cocktail with EDTA (Roche)). The nuclei were pelleted, washed once with complete lysis buffer, and then lysed with 100ul 0.5% SDS solution at 62°C with 400RPM agitation for <10 mins. 299 μl nuclease-free water and 20 μl of 20% Triton X-100 were added before 15 min incubation at 37 °C to quench SDS. 1X DpnII buffer (NEB), 0.8U/μl DpnII restriction enzyme (NEB), and 0.08 mg/μl BSA were supplemented to the chromatin solution for DpnII digestion at 37°C for 4 hours with 1000 RPM agitation. Additional 400U of DpnII was added to the sample for overnight digestion. DpnII was heat-inactivated at 62°C for <20 mins. An aliquot of the digested chromatin was sampled for DNA purification followed by DNA quantification, agarose gel electrophoresis, and PCR to monitor digestion efficiency. Digested chromatin equivalent to total 10 μg of DNA was ligated with 660U HC DNA ligase (Invitrogen, 30U/μl) in 1X ligation buffer (Invitrogen, supplemented with 1% Triton X-100, 0.1 mg/mL BSA) for 8-10 hours at 16°C without shaking. The reaction was incubated at room temperature for 30 mins. The chromatin was reverse-crosslinked with 0.5% SDS and 50 μg/mL of Proteinase K (Invitrogen) at 64°C overnight. RNA in the chromatin was digested with 2mg/mL RNase A at 37°C for one hour. The ligated DNA was extracted with phenol: chloroform: isoamyl alcohol (25:24:1) followed by chloroform and then precipitated with ethanol (68% to avoid SDS precipitation) in the presence of glycogen. The ligated DNA product was analyzed by agarose gel electrophoresis, and the DNA concentration was determined by Qubit DNA HS assay kit.

3 μg of the DNA product was further digested with 3U of specific second cutter (NlaIII, MseI, or MluCI). Restriction enzyme was heat-inactivated by incubating the chromatin at 65°C for 20 mins (MluCI was heat-inactivated in the presence of 1.17% SDS). The double-digested DNA was analyzed by agarose gel electrophoresis prior to phenol: chloroform: isoamyl alcohol (25:24:1) extraction and ethanol precipitation in the presence of glycogen. 1 μg of purified DNA was ligated with HC DNA ligase (Invitrogen, 30U/μl) at 20U/ug DNA in 1X ligation buffer (Invitrogen). For NlaIII or MulCI digested samples, the ligation was performed at 16°C overnight. For MseI digested samples, the ligation was performed at 4°C for 24 hours. The ligated DNA was recovered by phenol: chloroform: isoamyl alcohol (25:24:1) extraction and ethanol precipitation. 800 ng DNA from each of the biological triplicates were pooled to prepare 4C library (2.4 μg of DNA in total). The library for each viewpoint was constructed by PCR with 0.05 μM 4C adaptor primers (see primers section) and 2 ng/μl DNA templates using KAPA HiFi HotStart ReadyMix (KK2602). The libraries were subjected to size selection (250–500 bp) on a 4–20% TBE PAGE gel (Thermo Scientific). The recovered libraries were sequenced on the Illumina Nextseq platform using Nextseq 500 mid-output kit with 2×150 paired-end.

### Bioinformatics

The NGS data generated in this study have been deposited to the NCBI Gene Expression Omnibus (GEO) under accession number GSE141083.

For ChIP-seq analysis, reads were trimmed of adaptor sequences by TrimGalore ^4^ and were mapped by bowtie2 ^5^ against human reference genome GRCh38. The

PCR duplicates were removed by SAMtools rmdup ^6^. The mapping results were converted into signals by BEDTools ^7^ in bedGraph format. Then they were converted to bigWig format by bedGraphToBigWig ^8, 9^ for visualization on UCSC genome browser ^10^. Peak calling was performed by MACS2 ^11^.

RNA-seq reads were mapped by STAR ^12^ against human reference genome GRCh38 with GENCODE transcriptome annotation (v26) ^13^. RNA-seq reads were trimmed of adaptor sequences by TrimGalore ^4^ and then mapped by STAR to the human reference genome GRCh38 with reference gene annotation GENCODE 26 ^12, 13^. PCR duplicates were removed in the paired-end alignments by SAMtools rmdup. Alignments with mapping quality <20 were removed. Gene expression levels in FPKM were determined by cuffdiff ^14^ RNA-seq read counts are generated by featureCounts ^15^.

For RIP-seq, reads from the RIP data set and its control data set were trimmed of adaptor sequences by TrimGalore ^4^ and were mapped respectively by STAR ^12^ against the human reference genome GRCh38 with GENCODE ^13^ transcriptome annotation (v26). The resulting alignments were separated into two parts. (1) The exonic part consisted of alignments belonging to GENCODE annotated transcripts. (2) The non-exonic part consisted of the other alignments. Based on the number of reads that can be mapped into the transcriptome, the larger data set was shrunk down to fit the size of the smaller one. Then the normalized read count of each genomic position was generated for the EZH2 RIP data set and IgG control data set, respectively. The read coverage of each position was defined as the average of normalized read counts within ±150 bp region. Based on the comparison of the read coverage between the EZH2 RIP and the IgG control, sites with ≥ 2-fold enrichment and Poisson distribution p-value ≤ 10^−5^ were defined as peaks. Each peak was extended to surrounding areas until the fold enrichment dropped below 2. Peaks that can be found in both replicates within 1000bp range were merged and considered reproducible peaks. Sequences covered by reproducible peaks were extracted for further motif discovery analysis. The upstream and downstream sequences of reproducible peaks were also extracted and served as the background sequences in the motif finding analysis. Motif discovery is performed with the MEME suite using differential enrichment mode ^16, 17^.

For Nanopore sequencing, raw read collected from MinKNOW were converted to FASTQ file by Albacore 2.3.3 with the following arguments: -k SQK-RNA001 -f FLO-MIN106 -o fastq for direct RNA sequencing; or the following arguments: Albacore 2.3.1 with --flowcell FLO-MIN106 --kit SQK-PCS108 --output_format fastq for cDNA sequencing. For direct RNA sequencing, the reads were mapped to hg38 by Minimap2 ^18^ with -ax splice -uf -k14. The bam files were converted to bed12 by bamToBed -bed 12 and were converted to bigBed file by bedToBigBed -type=bed12. Only uniquely mapped reads were retained. For cDNA sequencing, the reads were mapped to hg38 by LAST with lastal -d90 -m50 -D10 and last-split -d2 ^19, 20^. The maf files were converted to psl file with maf-convert -j1e6 psl. The psl files were subsequently converted to bam file with psl2sam.pl and cigar_tweaker scripts within the SAMtools package ^6^. For visualization, the read alignments were converted into bedGraph by BEDTools ^12^, and finally, into bigWig format by bedGraphToBigWig ^8, 9^.

For 4C-seq analysis, the long-range genomic interaction regions generated by the 4C-Seq experiment were analyzed by the R package r3CSeq v1.30.0 ^21^. Briefly, for each replicate, the raw reads were aligned to the masked version of the reference human genome (masked for the gap, repetitive and ambiguous sequences) downloaded from the R Bioconductor repository (BSgenome.Hsapiens.UCSC.hg19.masked). The chromosome 11 was selected as the viewpoints, and DpnII was used as the restriction enzyme to digest the genome. A non-overlapping fragment was selected to identify the interaction regions. BAM files were converted to read coverage by bedtools genomecov. The read coverage was normalized according to the sequencing depth. The read coverage difference of the treatment and the control was calculated based on the normalized read coverage. The resulting bedGraph files were converted to bigWig format by bedGraphToBigWig. The genome version of bigWig files was then converted to hg38 by CrossMap ^22^. The profile of read coverage difference on genomic intervals was plotted by deepTools ^23^.

